# SARS-CoV-2 Spike Glycoprotein Receptor Binding Domain is Subject to Negative Selection with Predicted Positive Selection Mutations

**DOI:** 10.1101/2020.05.04.077842

**Authors:** You Li, Ye Wang, Yaping Qiu, Zhen Gong, Lei Deng, Min Pan, Huiping Yang, Jianan Xu, Li Yang, Jin Li

## Abstract

COVID-19 is a highly contagious disease caused by a novel coronavirus SARS-CoV-2. The interaction between SARS-CoV-2 spike protein and the host cell surface receptor ACE2 is responsible for mediating SARS-CoV-2 infection. By analyzing the spike-hACE2 interacting surface, we predicted many hot spot residues that make major contributions to the binding affinity. Mutations on most of these residues are likely to be deleterious, leading to less infectious virus strains that may suffer from negative selection. Meanwhile, several residues with mostly advantageous mutations have been predicted. It is more probable that mutations on these residues increase the transmission ability of the virus by enhancing spike-hACE2 interaction. So far, only a limited number of mutations has been reported in this region. However, the list of hot spot residues with predicted downstream effects from this study can still serve as a tracking list for SARS-CoV-2 evolution studies. Coincidentally, one advantageous mutation, p.476G>S, started to surge in the last couple of weeks based on the data submitted to the public domain, indicating that virus strains with increased transmission ability may have already spread.

## Introduction

On March 11^th^ 2020, the World Health Organization declared that the novel coronavirus disease (COVID-19) a pandemic crisis. The pathogen, namely severe acute respiratory syndrome coronavirus 2 (SARS-CoV-2), is the third coronavirus that causes serious global outbreak of the infectious diseases in 21^st^ century after Severe Acute Respiratory Syndrome Coronavirus (SARS-CoV) and Middle East Respiratory Syndrome Coronavirus (MERS-CoV). CoVs are by far the largest known positive-sense single-stranded RNA viruses with highly conserved genome in regions that encode the CoVs core proteins. SARS-CoV, MERS-CoV, and SARS-CoV-2 are members of the beta-coronavirus genus with the genome size of approximately 30 kilobases. CoV genome begins with a 5’ replicase gene that encodes the nonstructural proteins (Nsps), followed by four major virion structural protein coding genes that are crucial for the transmission and proliferation of the virus. Specifically, the spike glycoprotein (S, or spike) is mainly responsible for the cross-species and human-to-human transmission of CoVs via its interactions with the host cell receptors before inducing cell fusion. The small envelope protein (E) and the membrane protein (M) play important roles in viral assembly. Apart from its role as a structural protein, the nucleocapsid protein (N) also involves in the replication and transcription of the viral genomic information. The genome is capped at the 5’ with a 3’ poly(A) tail, similar to eukaryotic mRNA.

Comparative genomics research [1, 2] and studies with experimental validations [3, 4] have revealed the cell entry of SARS-CoV-2 depends on binding of the spike protein and the angiotensin-converting enzyme 2 (ACE2) receptor on host cells. Moreover, SARS-CoV-2 spike exhibits similar binding affinity to SARS-CoV spike protein with hACE2 [3], and the binding affinity correlates with the overall viral replication rate and the disease transmission rate [5-7], suggesting the comparable transmission rate between SARS-CoV-2 [8] and SARS-CoV.

During the epidemic outbreak of SARS-CoV in 2003, multiple genetically diverse SARS-CoV strains have been identified, but none of them showed significant higher virulence over others [9, 10]. According to recent studies and the phylogeny relationship of SARS-CoV-2 [11, 12], it is expected that the mutation rate of SARS-CoV-2 is similar to the majority of other RNA viruses including SARS-CoV [13, 14], with approximately 10^−6^ nucleotides per site per cell infection. As of April 12^th^ 2020, there are over 1.8 million infections spreading to more than 200 counties/areas for SARS-CoV-2 and the numbers are still escalating. In contrast, only ∼8,000 infections from 14 countries have been reported worldwide by the time SARS-CoV is contained. Consequently, with significantly higher number of infections in a pandemic scale, SARS-CoV-2 is expect to accumulate more mutations than SARS-CoV. Even though in theory, the virulence is expected to be high in the early stage of an epidemic, and will be decreased overtime until an endemic equilibrium is reached [15-19], there is still an urging need for studies on possible mutation outcomes on the transmission of the disease during COVID-19 pandemic.

In this study, we utilized existing models to identify key residues on SARS-CoV-2 spike protein that interacts with hACE2. We predicted possible amino acids substitutions that may significantly alter the binding affinity between spike and hACE2 using computational methods, thereby inferring the changes in the transmission rate of mutated virus strains. We then downloaded nearly 700 SARS-CoV-2 complete genome sequences from GISAID [20, 21] to identify potential strains embedding these high risk mutations. Results have shown that currently no mutations have been identified near the S-hACE2 interaction surface area. Although it seems that the whole spike receptor-binding domain (RBD) is subject to negative selection, there are still multiple predicted advantageous mutations near the S-hACE2 interaction surface area that could promote virus transmission. In fact, the number of strands with the advantageous mutations predicted by this study starts to stand out, indicating that virus strands with higher transmission may have already spread.

## Results and discussion

### Mutations in spike RBD and the predicted impacts on viral transmission

We inspected the hot spot residues that contribute to S-hACE2 interaction (figure 1a). Overall, 39 important residues within 6 Å around hACE2 were identified (table 1). Among these residues, 14 of them are directly interact with hACE2 via hydrophobic interaction or hydrogen bonding. We iterated all possible single nucleotide polymorphisms (SNPs) for each residue and analyzed the downstream impact of these amino acid alterations. Here, synonymous mutations and non-synonymous with ΔAffinity between -1.5 and 1.5 [22] were defined as neutral mutation, whereas non-synonymous mutation with ΔAffinity ≥ 1.5 or SNPs that lead to pre-termination stop codons are considered as deleterious. The rest of mutations are characterized as advantageous, since the S-hACE2 interaction are benefited from the amino acid substitution (ΔAffinity ≤ -1.5). For instance, figure 1b showed that the S-hACE2 binding could be benefit by a hydrogen bond formation between S^Lys498^ and hACE2^Gln42^, in addition to a salt bridge formation between S^Lys498^ and hACE2^Asp38^, despite a hydrogen bond loss between S^Gln498^ and hACE2^Gln42^ due to the p.498Q>K mutation. Comparatively, the p.498Q>E mutation can lead to a hydrogen bond formation between S^Glu498^ and S^Gly466^ that stabilizes the spike protein, which makes it less favorable to S-hACE2 interaction (figure 1c).

**Table 1.** Hot spot residues and their predicted mutation effects on S-hACE2 interaction. 38 hot spot residues are listed in the table with all mutations grouped by their predicted impact. Among these residues, several of them were less relevant to the spike-hACE2 binding. For instance, mutations on 474Gln, 479Pro, 490Phe and 492Leu were all predicted as neutral. Moreover, 487Asn seems to be a crucial residue because all non-synonymous mutations are deleterious. In contrast, none of the residues only contain advantageous mutations.

**Figure 1.**
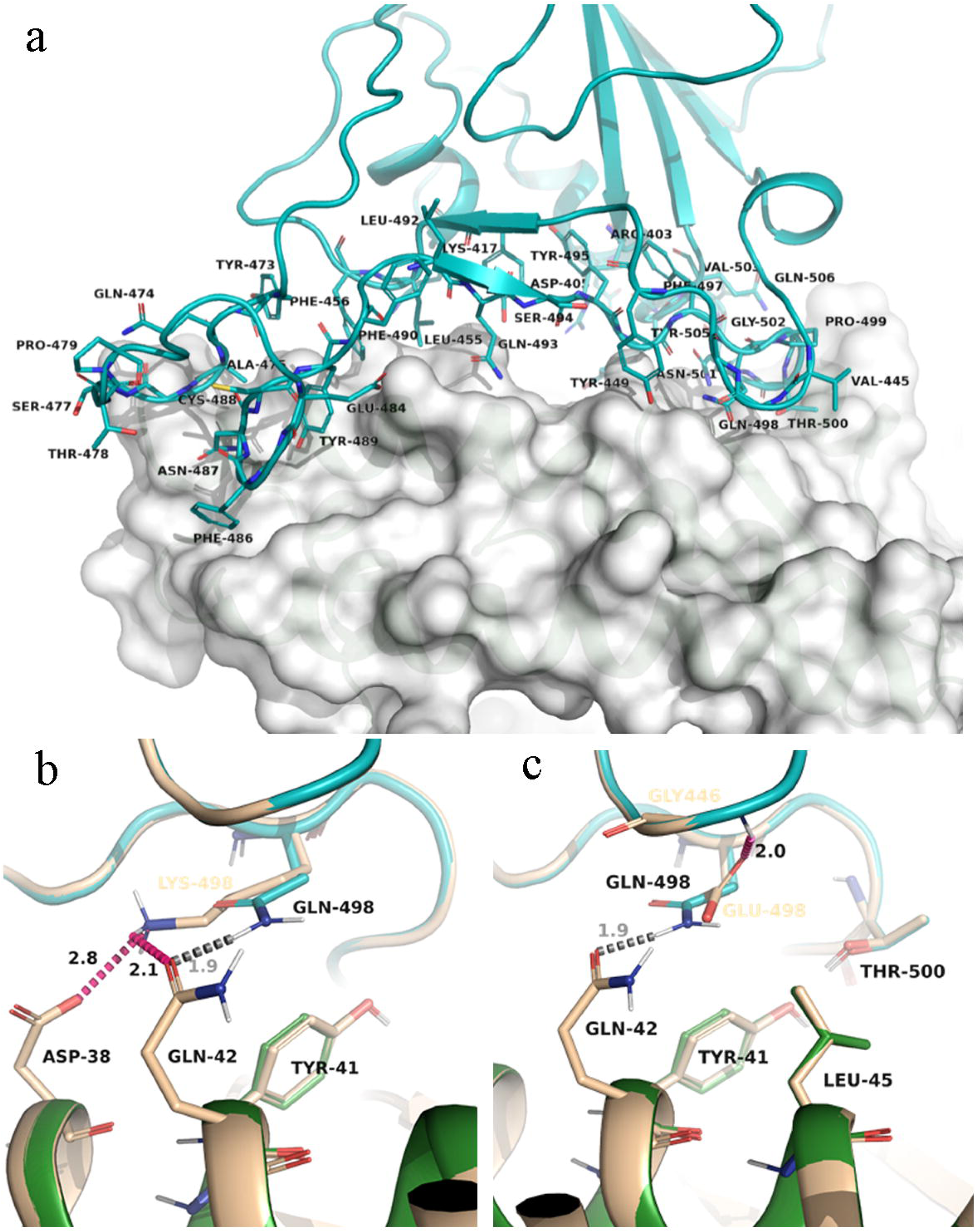
Hot spot residues and the impact of selected mutations to SARS-CoV-2 S-hACE2 interaction. SARS-CoV-2 S-hACE2 complex. (a) Residues from spike protein that are within 6 Å around hACE2 were labeled and adjusted for better visualization. Two mutations from the same residue has been chosen to illustrate the impact of the mutations to S-hACE2 interaction. (b) The p.498Q>K mutation breaks the hydrogen bond between S^Gln498^ and hACE2^Gln42^ (grey dash line) but forms a hydrogen bond between S^Lys498^ and hACE2^Gln42^, and a salt bridge between S^Lys498^ and hACE2^Asp38^, enhancing the binding interaction. (c) The p.498Q>E mutation results in the loss of the same hydrogen bond with hACE2^Gln42^ (grey dash line), in addition to the formation of a hydrogen bond within the spike receptor binding surface, which is unfavorable for S-hACE2 interaction.

### Low mutation frequencies in hot spot residues may be driven by host specificity selection

To scan SARS-CoV-2 mutations particularly for those occurring near the spike RBD, we downloaded all 696 SARS-CoV-2 genome sequences from GISAID at the time of writing this manuscript to check the number of mutations accumulated across the SARS-CoV-2 genome. Overall, we observed a dramatically low genetic diversity within the RBD of SARS-CoV-2 (figure 2). Moreover, no variants were found in the hot spot residues. This either suggested (1) the hot spot residues region is highly conserved during the evolutionary process, or (2) virus strains with reduced transmission abilities becomes minorities so that the mutations were not reported in the consensus genomic sequences. It is also possible that (3) unprecedented variants in hot spot residues might raise from the insufficient time of mutation accumulation.

**Figure 2.**
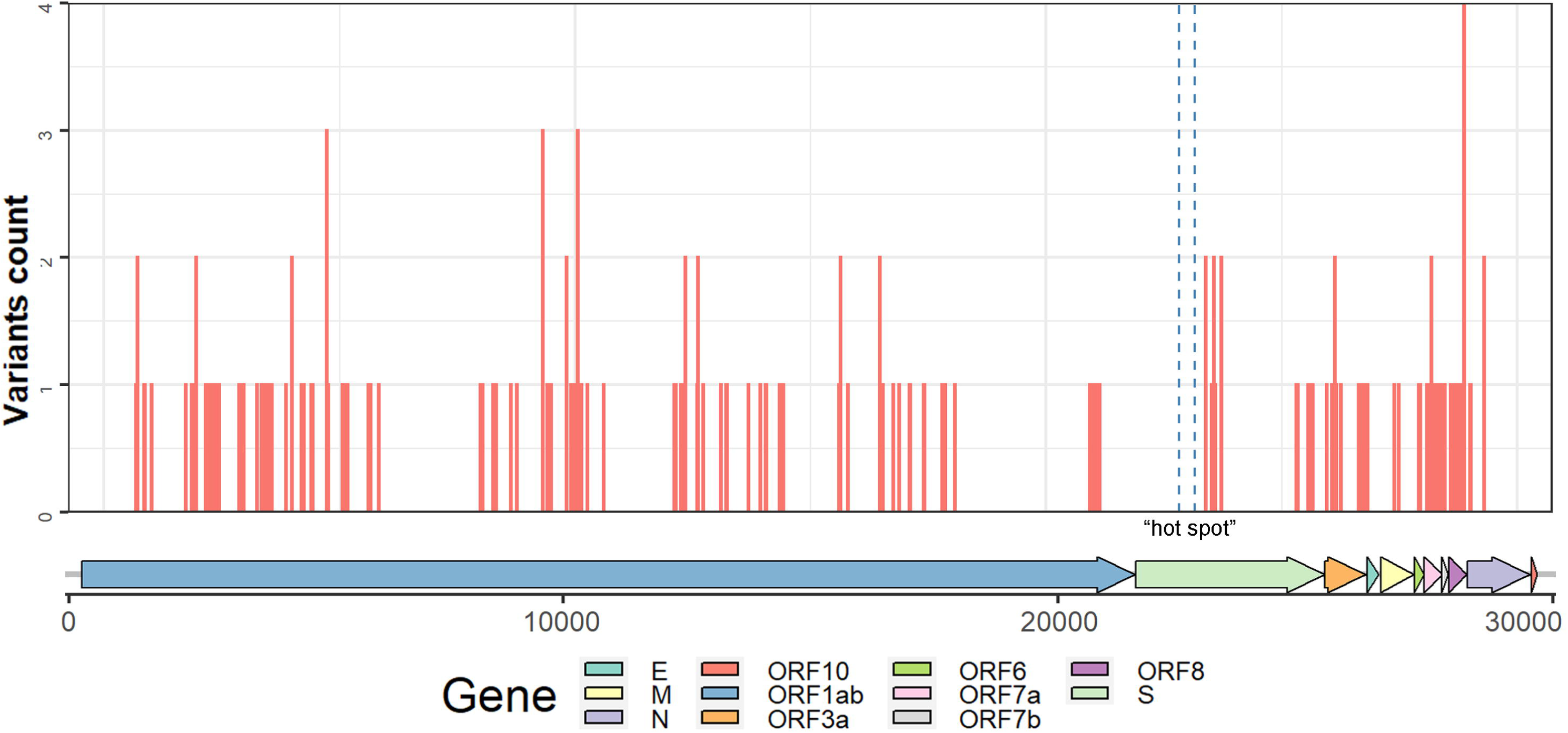
Genome wide SNP distribution. SNPs were called from multiple sequence alignment result and mapped to its genomic locations. 68% of the mutations are found in ORF1ab coding region. There are few mutations identified in spike protein, however, none of these mutations are in the hot spot region.

To verify (1) and (3), we compared the nucleotide diversity of the hot spot residue regions among SARS-CoV-2, SARS-CoV, and MERS-CoV, to see if mutations were reported in the hot spot residues from SARS-CoV-2 and MERS-CoV spike orthologs. The boundary of the hot spot residues for SARS-CoV and MERS-CoV were identified using the same approach of SARS-CoV-2 (supplementary file 1). As expected, we observed variants in both of the hot spot residues of SARS-CoV and MERS-CoV (supplementary file 2), which demonstrated the hot spot residues are not mutation-free. Additionally, we found the SARS-CoV-2 had an overall low nucleotide diversity (supplementary file 2) compared to SARS-CoV and MERS-CoV, and the MERS-CoV had the highest overall nucleotide diversity among the three viral species. These results further indicated that the relative short mutation accumulation time is one of the major reasons for the unprecedented variants in hot spot residue regions, even though COVID-19 has already been spread globally for several months.

As the key determinant of host specificity, spike protein comprises an N-terminal S1 subunit and a membrane-embedded C-terminal S2 region, in which the S1 domain was thought to be more variable than S2 domain normally [3, 23]. Therefore, the unprecedented variant in hot spot residues regions were likely suffered the host specificity selection, allowing the stabilized transmissions within species. However, despite of the negative Tajima’s D in the hot spot residues regions of SARS-CoV and MERS-CoV (supplementary file 2), we still cannot exclude the possibility of positive selection, as the negative Tajima’s D is more likely caused by the expansion of virus population. Collectively, we didn’t find any mutations in the hot spot residues region in SARS-CoV-2 in our study, which might be driven by the host specificity selection.

### Selection pressure analysis on spike and spike RBD

In spite of the crucial role of host specificity selection in maintaining the transmission capacity of viruses, adaptive evolution still ubiquitously happened on viral genomes during animal-to-human transmission and human-to-human transmission [24, 25]. We analyzed the ratios of the rates of non-synonymous to synonymous changes (dN/dS) [26] for three coronaviruses with a focus on spike protein and spike RBD. We detected positive selection in the spike gene of SARS-CoV-2 and purifying selection for both SARS-CoV and MERS-CoV, although only MERS-CoV purifying selection was statistically significant (p-value<0.001, figure 3). Such observation correlated with the selection pressure analysis on SARS-CoV spike protein during different epidemic phases [10] and overall dN/dS ratio reported for MERS-CoV spike protein [27]. We further inspected the scaled-dN/dS ratio in spike functional domains, particularly RBD. MERS-CoV and SARS-CoV spike RBD were both under purifying selection, even though multiple positive selection sites on MERS-CoV spike were identified during the MERS epidemic [28, 29]. In contrast, the positive scaled-dN/dS value of SARS-CoV-2 spike protein was largely contributed by the region between RBD and S2 domain. In fact, the scaled-dN/dS for SARS-CoV-2 RBD was below zero, suggesting that overall the RBD was not under positive selection.

**Figure 3.**
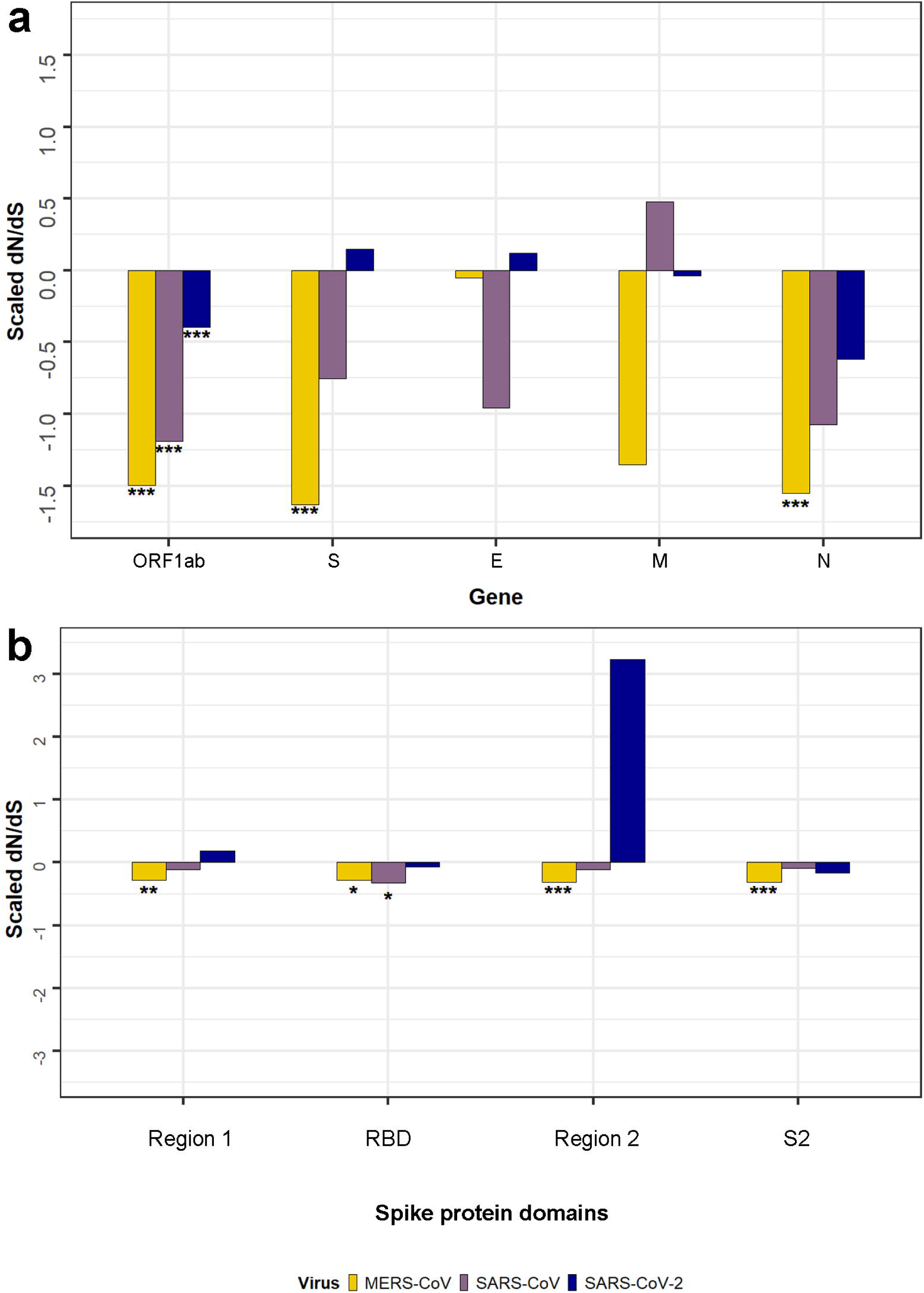
Comparison of dN/dS for major genes and spike protein domains of SARS-CoV-2, MERS-CoV, and SARS-CoV. (a) The scaled dN/dS value of five major genes. ORF1ab and N were under purifying selection in all three species. For spike protein, only SARS-CoV-2 spike was under positive selection. (b) However, the positive scaled-dN/dS value was mainly contributed by two non-domain regions, that is region 1, which is on the 5’ end of RBD, and region 2, which is between RBD and S2 domain. Due to limited number of mutations identified so far, none of the scaled-dN/dS values for these regions/domains were statistically significant. It is worth mentioning that the spike RBD for MERS-CoV and SARS-CoV were under purifying selection with statistical significance, suggesting the coronavirus spike RBD are usually under purifying selection eventually.

### The hot spot residues are more likely to be deleterious when mutation happens

Hu et al. has conducted a genome-wide evolution and variation analysis using 42 complete genome sequences of the SARS-CoV [30]. These include all 32 complete genome sequences of the SARS-CoV collected from GenBank as of Aug 17^th^, 2003, two months after the end of the SARS epidemic, along with 10 genomes sequenced by Beijing Genomics Institute. We believe these sequences are enough to reflect the mutation occurrences of SARS-CoV during the emergent pneumonia epidemic in 2003. We utilized the SARS-CoV transition and transversion statistics (reformatted in supplementary file 3) from this study to infer the probabilities of amino acid changes for each of the 38 residues listed in table 1, given the homogeneity of SARS-CoV-2 and SARS-CoV. A complete list of mutations and their probabilities are shown in table S1 (supplementary file 4). Taking Gln498 as an example, based on transition and transversion rate at each codon phase from SARS-CoV, we estimated that when a mutation occurs at SARS-CoV-2 S^Gln498^, there is a 15.4% chance to be synonymous, 33.3% chance to be advantageous, 0% chance to be non-synonymous neutral mutations, and 51.3% chance (with a 17.9% chance of a pre-terminated stop codon mutation) to be deleterious. We highlighted 19 residues in figure 4 with higher chances to develop functional mutations (p_advantageous_+p_deleterious_ ≥ p_neutral_) when non-synonymous mutation happens. It is obvious that the majority of these high impact mutations are deleterious, suggesting that random mutations of these amino acids are more likely to decrease the S-hACE2 binding affinity, thereby reducing the transmission ability of the mutated strain. Such mutations may not be detectable by sequencing or are likely to be classified as rare mutations that are often excluded in the consensus genome sequences.

**Figure 4.**
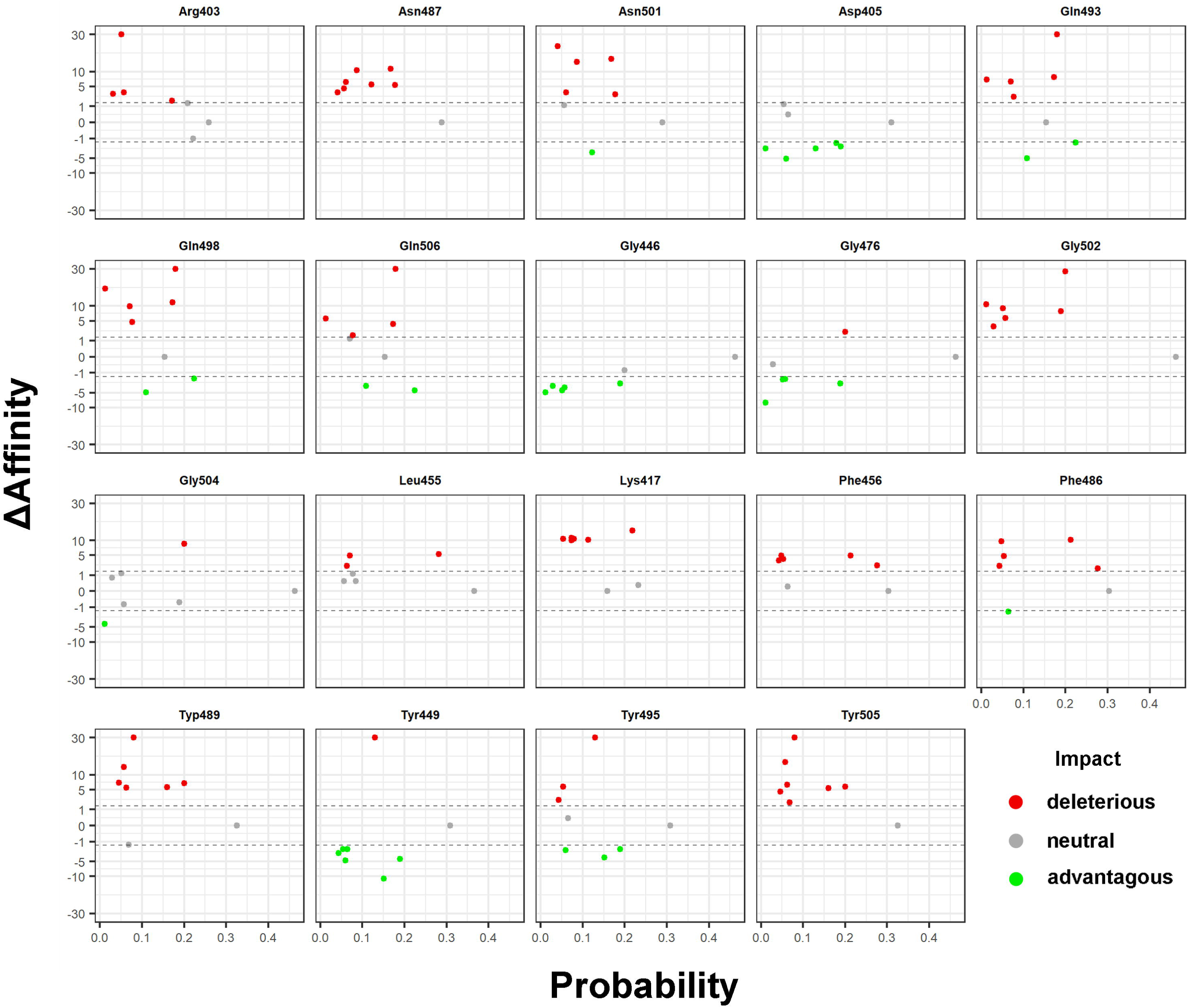
Functional mutation probabilities and the impact of the mutations of 19 selected residues. Residues from the hot spot residues were selected if they are more likely to develop a functional impact on SARS-CoV-2 S-hACE2 interaction. Possible mutations of each residue is plotted in scatter plot separately, with each dot represent a unique amino acid substitution, colored by its downstream impact. For each dot, the x-axis is the predicted mutation probability and the y-axis is the Δaffinity in log-scale. Overall, the number of red dots (deleterious mutations) are higher than the green (advantageous mutations) and grey (neutral mutations) dots, suggesting that these mutations are more likely to have deleterious mutations. However, dots from Asp405, Gly446, Tyr449, Gly476, Tyr495 and Gln506 were mostly green, indicating higher chances of developing advantageous mutations when these residues mutated.

Previously multiple positive selection sites on MERS-CoV spike proteins were reported during its epidemic. Comparatively, only a limited number of mutations were identified near the hot spot residue region for SARS-CoV-2. Therefore, it is challenging to apply the same methods on SARS-CoV-2. Alternatively, we analyzed possible mutations of the hot spot residues and predicted multiple amino acids that may under strong positive selection. Specifically, Asp405, Gly446, Tyr449, Gly476, Tyr495, Gln506 are 6 residues with p_advantageous_ > p_deleterious_. These hot spot residues require closer surveillance, as mutations surged from these sites could indicate mutated strains with higher transmission abilities.

### Analyze hot spot residue mutations from public database

The number of confirmed COVID-19 cases grows exponentially as we conducted this study, with 10 times more SARS-CoV-2 genome sequences submitted to GISAID in a month, reaching over 6,000 sequences as of Apr. 12^th^, 2020. Instead of collecting the complete list of publically available sequences that are expected to grow each day, we chose to verify our prediction with the mutation database from CNCB (the China National Center for Bioinformation, https://bigd.big.ac.cn/ncov/) directly. Among 2,645 mutations, only 22 SNPs were mapped to the hot spot residue region with a genomic coordinates between 22,769 and 23,080, six of which were synonymous mutations (c.1518caA>caR was wrongly annotated as non-synonymous mutation. See supplementary file 5). Apart from the mutations that directly fell into hot spot residues, we also evaluated the ΔAffinity value for all amino acid substitutions presented in hot spot residue region. Figure 5 illustrated that mutations with high number of strands were either neutral mutations (p.475Q and p.483V>A, with six and 16 strands, respectively), or advantageous mutations that could potentially increase the virus transmission (p.476G>S, eight strands). Although the number of strands with the mutation is not necessarily related to infectivity directly under the pandemic situation, the fact that variants with higher frequencies were not deleterious could imply that virus strands with deleterious mutations may exhibit restricted infectivity. On the other hand, the advantageous p.476G>S mutation was associated with the second highest number of strands in this region. Whether the high number is caused by oversampling from the same spread chain or by multiple strains evolved with the same positively selected strains need further investigation, even though the latter one seems more probable because incidences were reported from a couple of geographically distant countries (Belgium and USA) by different institutes.

**Figure 5.**
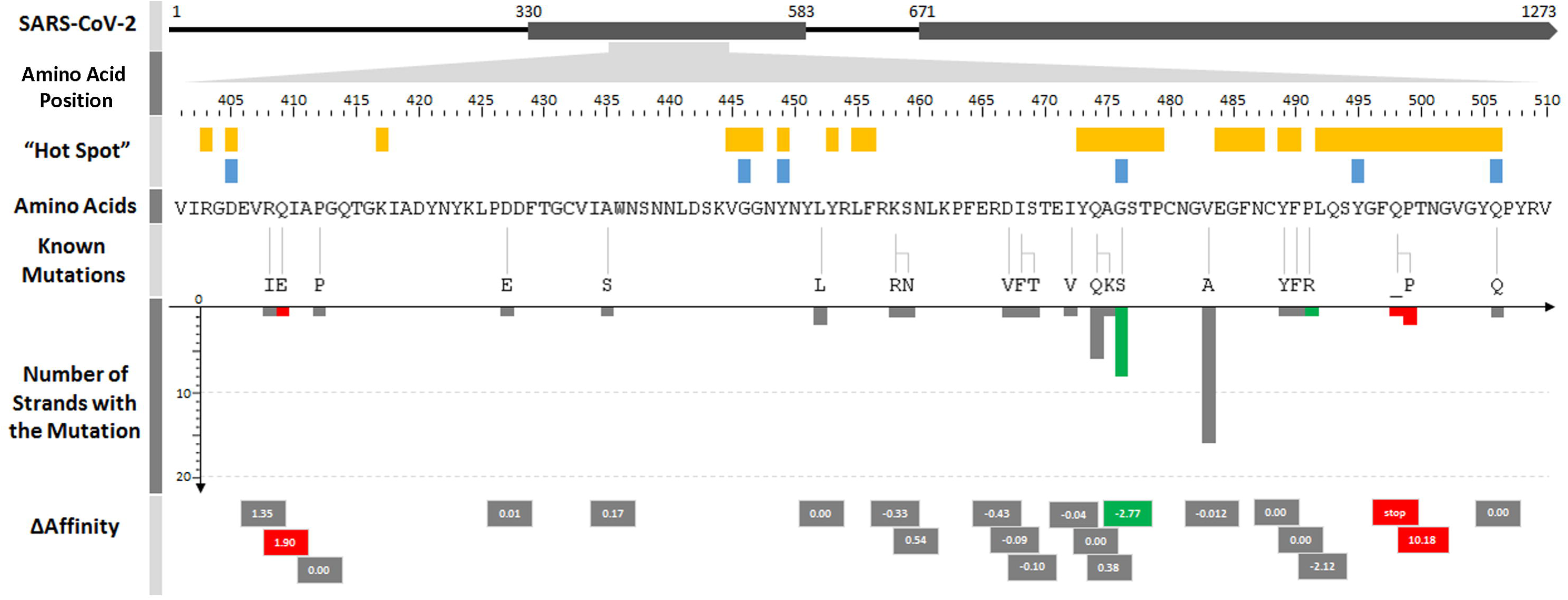
Mutation distribution on hot spot residue region using latest mutation statistics. Hot spot residues (yellow blocks) and six residues mentioned in figure 4 (blue blocks) were mapped to the amino acid positions of the spike protein. The wild type amino acid sequences and the mutations identified in this region are aligned together in Amino Acids and Known Mutations track. For each mutation, the number of strands with the mutation is plotted in the bar chart track. Boxes with numbers in the last track can be projected to the x-axis to track the Δaffinity of the corresponding mutations. Deleterious, neutral, and advantageous mutations are filled with red, grey and blue in the last two tracks, respective.

## Material and methods

### Hot spot residues identification and mutation analysis

The structure of the SARS-CoV-2 spike with hACE2 complex was initially downloaded from China National Microbiology Data Center (http://nmdc.cn/nCoV) with an accession number NMDCS0000001 (PDB accession: 6LZG). The analysis was performed using PyMol [31] (version 2.3.0) and Residue Scanning module of BioLuminate [32, 33] from Schrödinger suite. Residues on spike receptor binding domain (RBD) within 6 Å around hACE2 were considered as hot spot residues. The hot spot residues for SARS-CoV S-hACE2 (PDB accession: 6CS2) and MERS-CoV S-hDPP4 (PDB accession: 4L72) were identified using the same approach. For SARS-CoV S-hACE2 analysis, the side-chain prediction with 4.5 Å Cbeta sampling was selected to perform the side-chain prediction including sampling of the CA-CB orientation. With one single nucleotide mutation at a time, all possible amino acid substitutions for each of the hot spot residues were iteratively introduced to the wild type protein. Residue Scanning of BioLuminate systematically evaluates the binding affinity of residue mutations in the protein-protein interface. ΔAffinity parameter was used as indicators for binding affinity changes of the mutant protein as well as any other specific chains. Specifically, positive ΔAffinity values associate with reduced binding affinity of the mutant spike protein and vice versa. Therefore, amino acid substitution with ΔAffinity≥1.5 [22, 34] were defined as deleterious because it weakens the S-hACE2 interaction and may lead to reduced transmission rate of the coronavirus. Similarly, substitutions with Δ Affinity ≤ -1.5 were considered as advantageous, because enhanced binding may lead to higher viral transmission rate. Substitutions with ΔAffinity between -1.5 and 1.5 were labeled as neutral mutations, since the affinity changes seems to be negligible computationally. We also assigned Δ Affinity score of 0 for synonymous mutations and ΔAffinity score of 30 for pre-terminated stop codons for ease of illustration.

### Sequence variations and evolution analysis

The complete genome sequences for 696 SARS-CoV-2, 260 SARS-CoV, and 479 MERS-CoV (supplementary file 6) were used to conduct the population genetics and evolutionary analyses. Sequences for SARS-CoV-2 were downloaded directly from GISAID database. Sequences for SARS-CoV and MERS-CoV were queried from NCBI virus database [35] using the *blastn* function implemented in BLAST V2.6.0 [36] with their reference genomes (SARS-CoV: NC_004718.3; MERS-CoV: NC_019843.3), respectively. For sequence variation and evolution analysis, we firstly performed multiple alignment at genome levels within each species using MAFFT V7.455 [37]. The alignments were manually inspected and truncated to make sure the total lengths were consistent with the corresponding reference genome. The nucleotide diversity and Tajimas’ D value for each virus were calculated using the PopGenome V2.7.5 package [38] in R. A sliding window approach (window size = 100 bp, jump = 100 bp) was used to this calculation. We then calculated the dN/dS ratio using SNPGenie [39]. The gene information for the three viruses were extracted from the corresponding GFF files available in NCBI. The domain information of SARS-CoV-2 spike protein (NCBI accession ID: YP_009724390.1) was determined using Pfam [40] via sequence search.

### SARS-CoV-2 mutation probabilities

The occurrences of transition and transversion statistics from this study [30] were used to calculate per-nucleotide mutation at each codon phase. For instance, SARS-CoV-2 S^Gln498^ is encoded by a ‘CAA’ triplet. Using IUPAC notation [41], the probability of ‘CA<u>A</u>’ mutated to ‘CA<u>G</u>’ can be derived from the equation below, 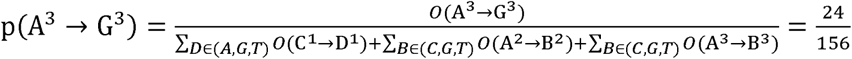 Here *O*(A^3^ → G^3^) stands for the occurrences of A>G mutation at the third codon phase. Therefore, we can conclude that if a mutation at S^Gln498^ happens, 15.385% it will be a synonymous mutation since ‘CAA’ and ‘CAG’ both encode glutamine.

### Hot spot residue mutation check

We downloaded a complete list of mutations for SARS-CoV-2 from CNCB on Apr. 12^th^, 2020. 22 single point mutations between 22,769 and 23,080 were extracted. Nucleotide substitutions described using IUPAC nomenclature without accurate annotation were manually curated. Δ Affinity of these 22 mutations were also calculated using BioLuminate if not available. The list of all mutations can be found in supplementary file 5.

## Authors’ contributions

YL conceived the study. YL and YW performed the bioinformatics analysis. YQ and ZG performed the protein-protein interaction analysis. LD collected the data. MP, HY, JX provided input for the study. YL and YW wrote the manuscript with input from all authors. LY and JL supervised the study and revised the manuscript.

## Acknowledgements

We gratefully acknowledge the authors, originating and submitting laboratories of the sequences from GISAID’s EpiFlu(tm) Database on which this research is based.

## Supplementary material

### Supplementary file 1

Hot spot mutation region for SARS-CoV spike-hACE2 (page 1) and MERS-CoV spike-hDPP4 (page 2). Residues witin 6Å around the cell surface receptor is also listed.

### Supplementary file 2

Population diversity distribution for SARS-CoV-2, SARS-CoV, and MERS-CoV genomes. The population diversity is evaluated by nucleotide diversity (upper track) and Tajima’s D (lower track) using a sliding window of 100bp. Hot spot regions of all three coronaviruses exhibit low population diversity.

### Supplementary file 3

Mutation statistics for SARS-CoV from Hu et al. The mutation count of X to Y is used for Y to X as well.

### Supplementary file 4

Mutation frequencies and the Δaffinity scores of all possible mutations on hot spot residues. Mutation type was set to synonymous and non-synonymous (non-synonymous mutation with stop codon was explicitly labeled). The impact of the mutation was set to advantageous, neutral and deleterious.Δaffinity scores of zero and 30 were set to synonymous mutations and pre-terminated stop codon mutations, respectively.

### Supplementary file 5

A list of all mutations downloaded from CNCB as of Apr. 12^th^, 2020. Mutations between 22,769 and 23,080 was highlighted in the table.

### Supplementary file 6

Accession IDs of all sequences used for this study. For SARS-CoV-2 genome sequences, a complete list of acknowledgement to all submitters was included in this file.

