## Supplementary file 1 for "SARS-CoV-2 Spike Glycoprotein Receptor Binding Domain is Subject to Negative Selection with Predicted Positive Selection Mutations"

### Slide 1
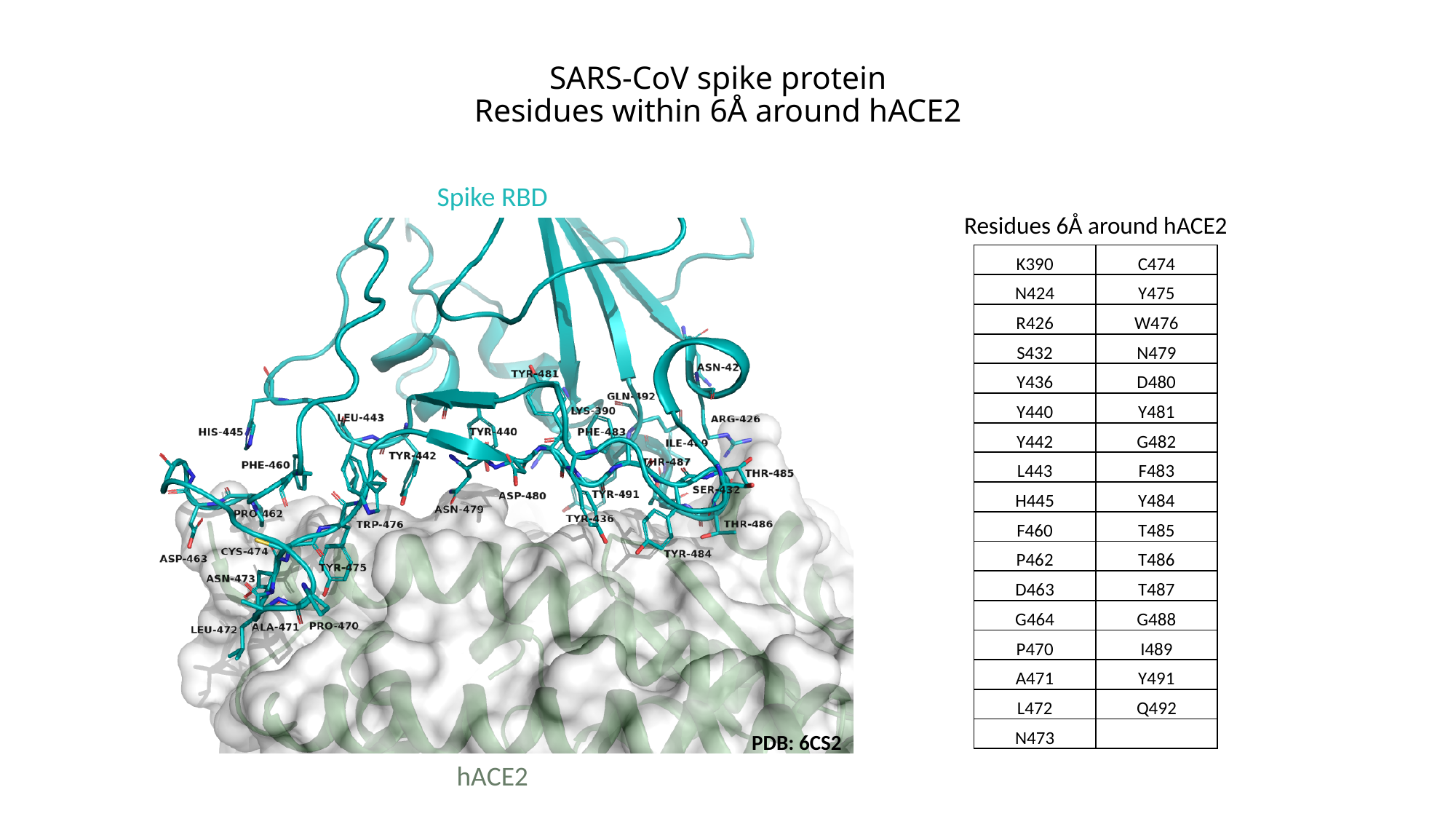

SARS-CoV spike protein
Residues within 6Å around hACE2
Spike RBD
PDB: 6CS2
hACE2
Residues 6Å around hACE2
| K390 | C474 |
| --- | --- |
| N424 | Y475 |
| R426 | W476 |
| S432 | N479 |
| Y436 | D480 |
| Y440 | Y481 |
| Y442 | G482 |
| L443 | F483 |
| H445 | Y484 |
| F460 | T485 |
| P462 | T486 |
| D463 | T487 |
| G464 | G488 |
| P470 | I489 |
| A471 | Y491 |
| L472 | Q492 |
| N473 | |

### Slide 2
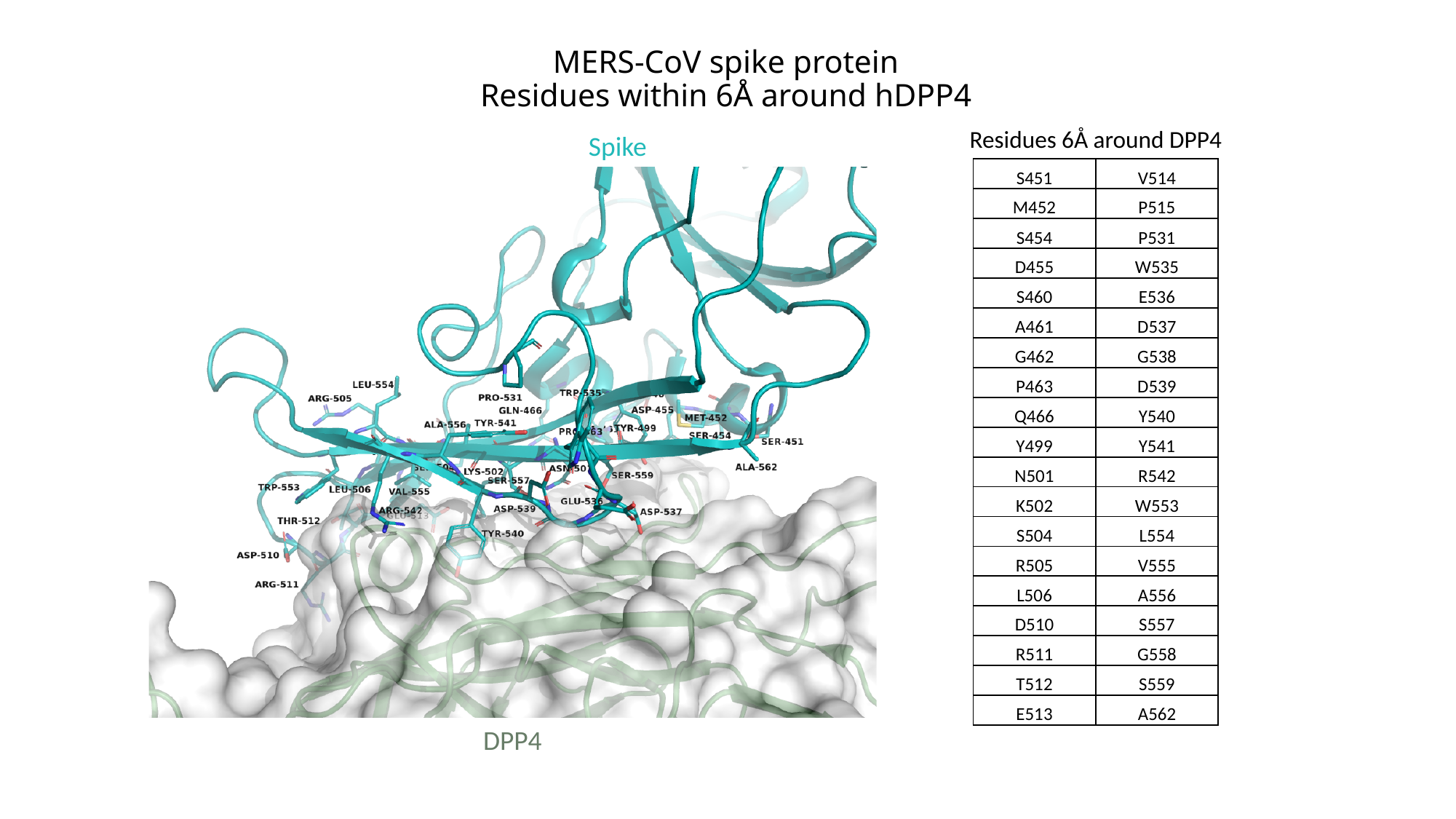

MERS-CoV spike protein
Residues within 6Å around hDPP4
Residues 6Å around DPP4
Spike
| S451 | V514 |
| --- | --- |
| M452 | P515 |
| S454 | P531 |
| D455 | W535 |
| S460 | E536 |
| A461 | D537 |
| G462 | G538 |
| P463 | D539 |
| Q466 | Y540 |
| Y499 | Y541 |
| N501 | R542 |
| K502 | W553 |
| S504 | L554 |
| R505 | V555 |
| L506 | A556 |
| D510 | S557 |
| R511 | G558 |
| T512 | S559 |
| E513 | A562 |
DPP4
