## Supplementary figures and images for "SARS-CoV-2 Spike Glycoprotein Receptor Binding Domain is Subject to Negative Selection with Predicted Positive Selection Mutations"

### Supplementary file 2

#### Slide 1
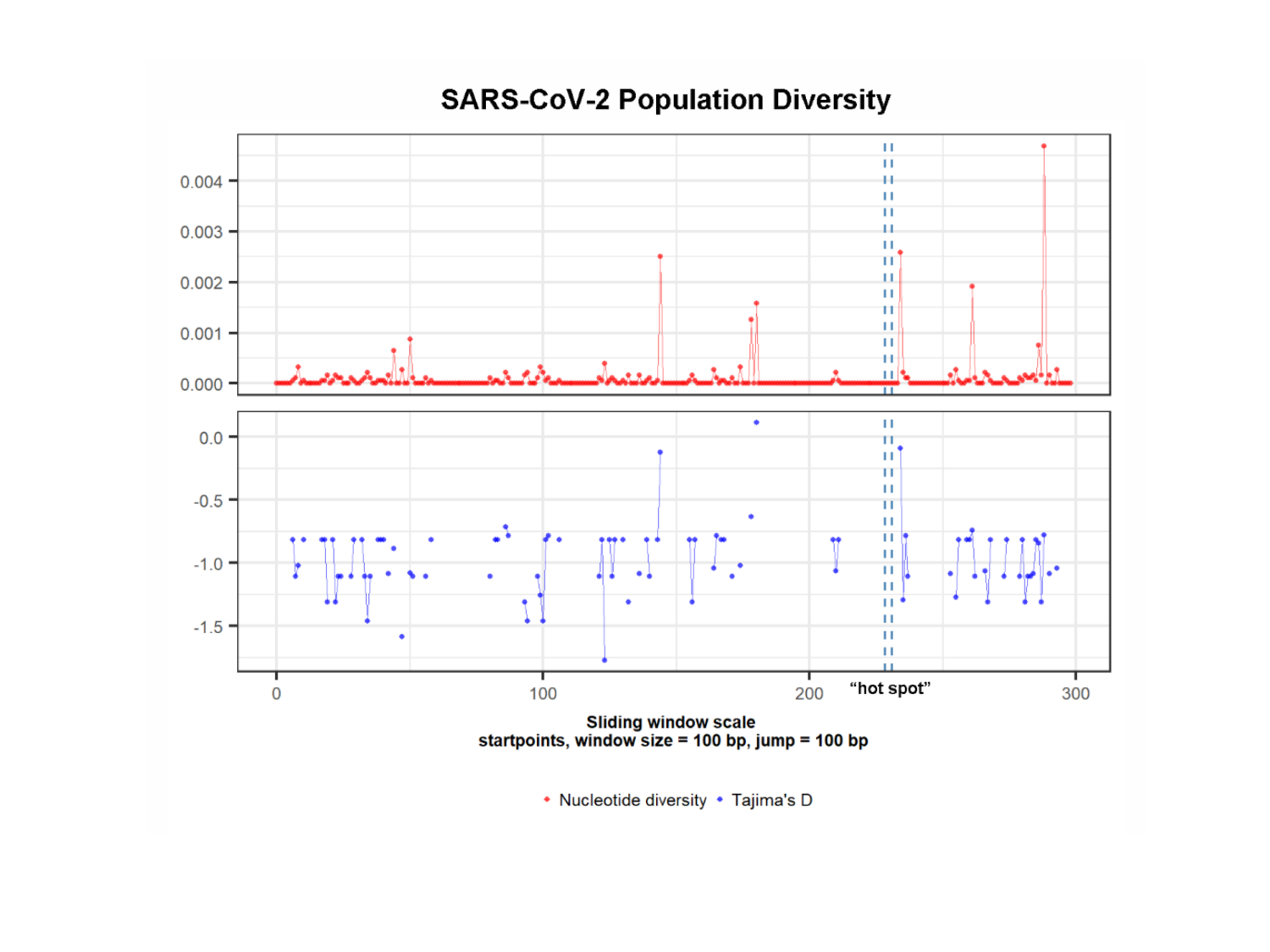

#### Slide 2
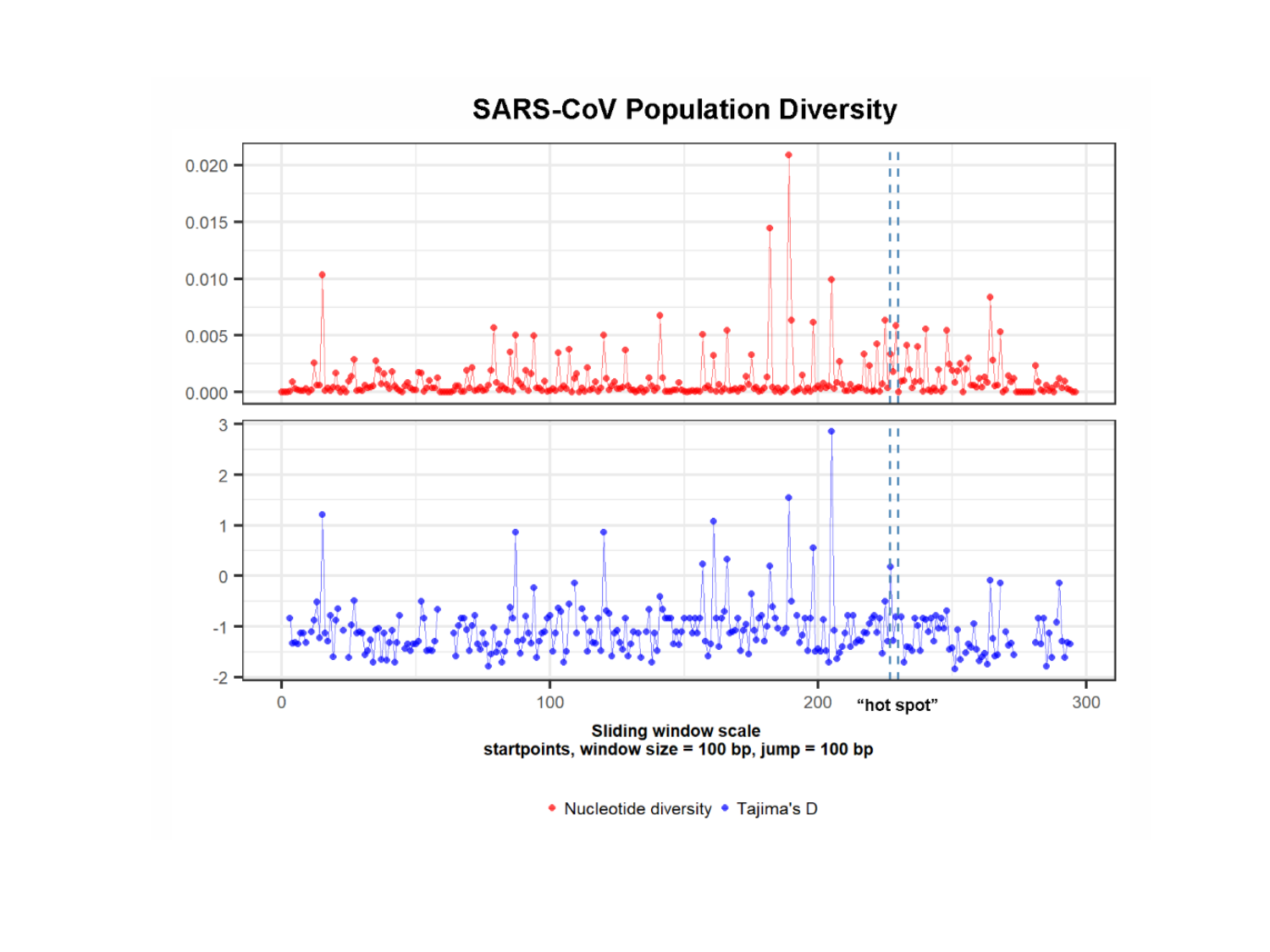

#### Slide 3
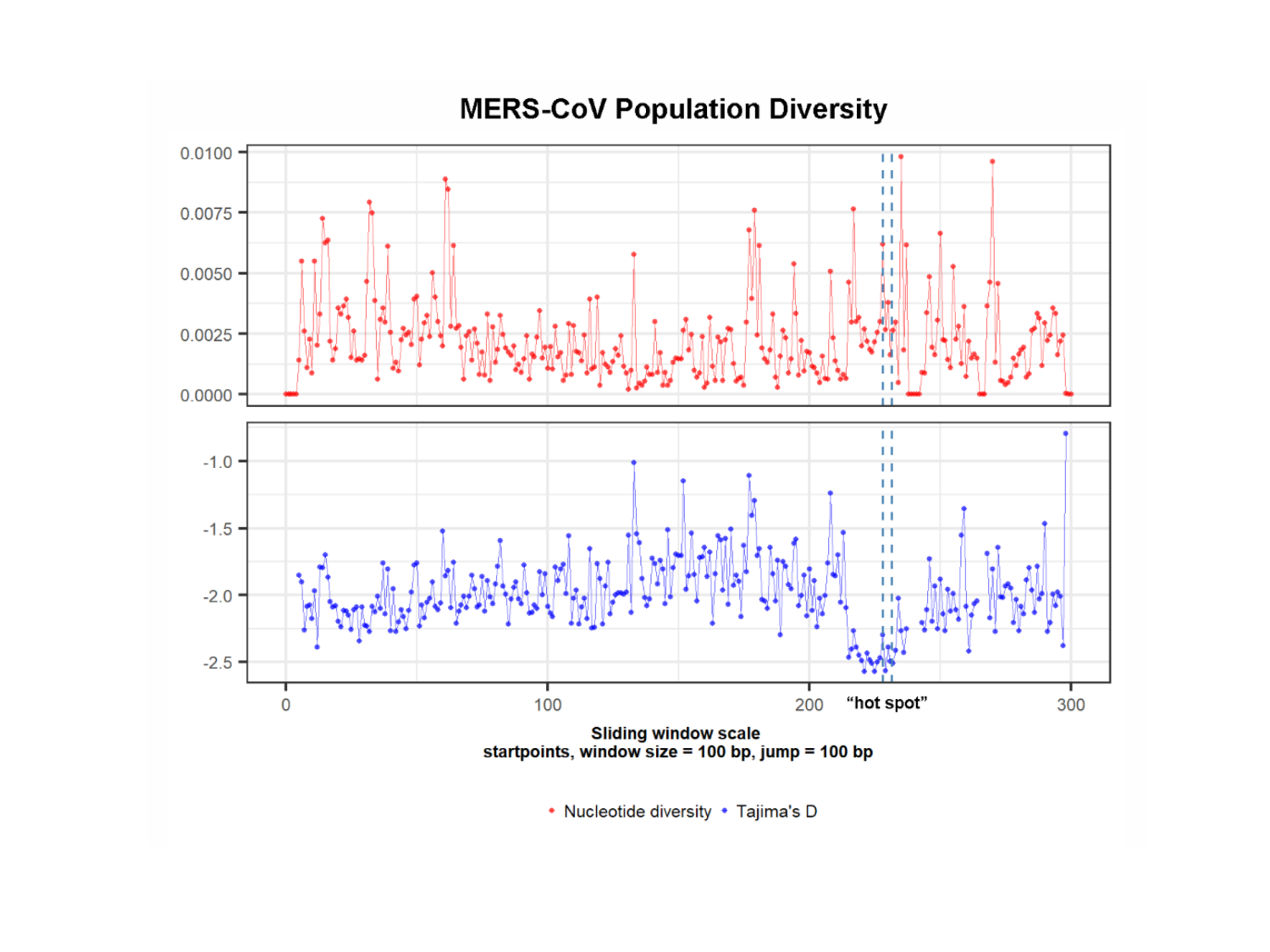
